# 2.5D Tractions in monocytes reveal mesoscale mechanics of podosomes during substrate indenting cell protrusion

**DOI:** 10.1101/2021.06.18.449040

**Authors:** H Schürmann, A Russo, AD Hofemeier, M Brandt, J Roth, T Vogl, T Betz

## Abstract

Degradation and protrusion are key to cellular barrier breaching in cancer metastasis and leukocyte extravasation. Cancerous invadopodia and myelomonocytic podosomes are widely considered as structural tools facilitating these processes and are thus summarized under the term invadosomes. Despite similar behaviour on the individual scale, substantial differences have been reported to arise on the collective scale. They are considered to be a result of podosome mesoscale-connectivity. In this study, we investigated global in-plane and out-of-plane mechanical forces of podosome clusters in ER-Hoxb8 cell derived monocytes. We are able to correlate these forces with the interpodosomal connectivity. The observed traction and protrusion patterns fail to be explained by summation of single podosome mechanics. Instead, they appear to originate from superimposed mesoscale effects. Based on mechanistic and morphological similarities with epithelial monolayer mechanics, we propose a spatiotemporal model of podosome cluster mechanics capable of relating single to collective podosome mechanical behaviour. Our results suggest that network contraction-driven (in-plane) tractions lead to a buckling instability that contributes to the out-of-plane indentation into the substrate. First assigning an active mechanical role to the dorsal podosome actomyosin network, we aim at translating actomyosin hierarchy into scale dependency of podosome mechanics.

## Introduction

Breaching of physical barriers (Fig. 1A) is an essential step in leukocyte extravasation ^1^ and cancer metastasis ^2^. Podosomes in healthy cells and invadopodia in invasive cancer cells that intrinsically combine matrix metalloproteinase (MMP) mediated degradation ^3,4^ and F-actin driven protrusion ^5,6^ have been associated to these processes ^7^. Despite functional similarities, their opposing biological roles make a tight distinction crucial for understanding and potential drug targeting. Over the past two decades, various differences have been highlighted. Most important here are differences in podosome cluster organisation and the collective dynamical and degradational behavior^8,9^. In recent years understanding of single podosome behaviour advanced greatly and by assigning mechanical and mechanosensory activity to these structures, mechanobiology was introduced into the field ^10,11^. Although podosome interconnectivity was linked to mesoscale dynamics ^8,12^, details on the link between the mechanical connectivity and single podosome mechanics remain largely unknown. Likewise, possible collective, hence global effects on the generation of cluster wide protrusions have hardly been addressed.

**Fig. 1.**
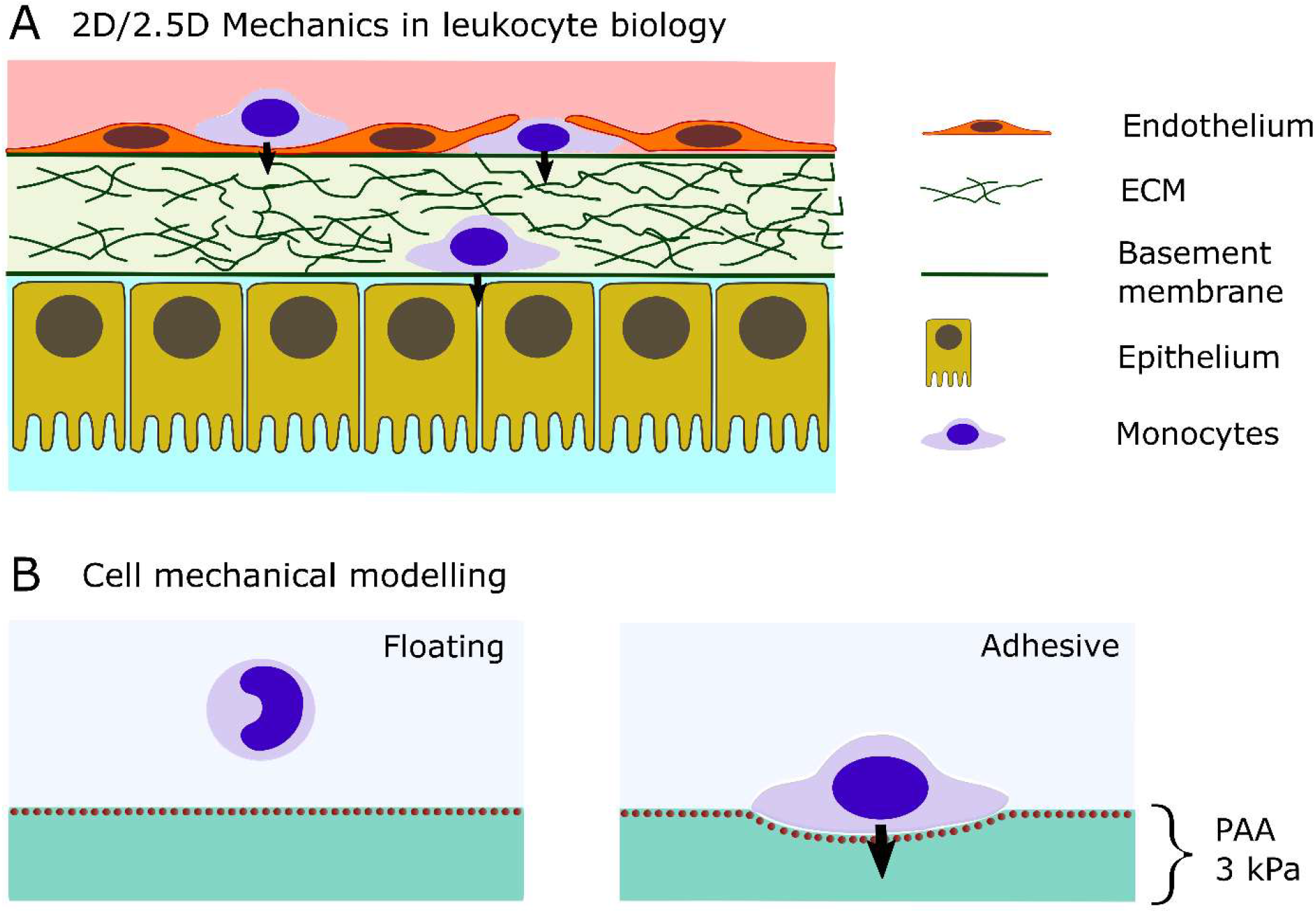
2D/2.5D Environments in leukocyte biology and cell mechanics. **(A)** Immune cells are frequently required to move out-of-plane at approximated 2D surfaces. The surfaces involved can be composed of the vascular endothelial cell lining or endothelial/epithelial basement membranes, respectively. **(B)** The mechanical aspects of these 2D/2.5D situations can be modelled by fibronectin coated soft polyacrylamide (PAA) hydrogels. Embedded marker beads, electrostatically accumulated close to the surface, allow to deduce the in-plane and out-of-plane deformations from which 3D tractions can be calculated.

Podosomes are known to form complex, hierarchical actin networks. These result in various (super)structures that reach far beyond the characteristic punctate actin protrusions. Among the most frequently investigated are podosome clusters. While at lower resolution the fluorescence obtained from F-actin generates an interpodosomal ‘cloud’ ^9^, at higher resolution its fine-tuned nano-architecture becomes apparent. Here, an architectural correlate of mesoscale-connectivity could be identified in the connecting cable network that spans between adjacent podosomes ^13^. Although connectivity was previously assumed to be sufficiently considered as a passive mechanical coupling ^12^, more recently a myosin II dependent dynamical role suggests mechanical activity ^8^.

Non-muscle myosin II is frequently coupled to F-actin strands or higher-order networks thus coining the term actomyosin. With its capability to generate contractile forces via actin translocation it was shown to play a major role in cellular force generation ^14,15^ thus implying mechanical activity where present. Regarding the ventral (i.e. substrate faced) podosome actin network, three major elements can be distinguished: the characteristic F-actin rich core as well as the myosin II decorated F-actin based lateral fibers and connecting cables ^13^. As assumed by composition, for myosin II decorated lateral fibers a role in protrusion force generation could be identified on the single podosome level ^16^. In contrast, a corresponding active mechanical role for the densely myosin II decorated connecting cable network has yet to be recognized.

A putative connectivity associated effect was reported for degradation assay studies: Individually, podosomes like invadopodia locally degrade and protrude into the substrate. On the cell scale, individual invadopodia still dominate ^17^. Opposingly, podosomes do not degrade and indent focally and deeply but homogeneously and shallowly upon cluster formation ^18^ suggesting altered mechanics on the collective scale for podosomes.

In this study, we explored 3D tractions underlying protrusion exerted by podosome cluster bearing cells on a compliant 2D hydrogel (Fig. 1B) by Traction Force Microscopy ^19^ (TFM). Compared to most previous studies we here access the vertical force along with planar forces. This approach is typically referred to as 2.5D TFM. To minimize biological variance, we employed murine Lifeact-EGFP labelled ER-Hoxb8 derived monocytic cells. This cell model poses a recently established estradiol-dependent, conditionally immortalized system of myeloid progenitor cells ^20,21^. Previously, these cells were successfully used in cell migration studies and have been shown to be comparable to standard primary cells regarding podosome formation and degradation ^22^.

We demonstrate that differentiated ER-Hoxb8 monocytes regularly display podosome clusters upon treatment with S100A8/A9, an endogenous protein secreted in systemic inflammation and also known as calprotectin ^23,24^, as well as *lipopolysaccharide* (LPS), an endotoxin of gram-negative bacteria capable of triggering inflammation ^25^. Followed by traction and stress analyses we resolve a cluster-wide, two-component traction pattern with no mutual correlation to the punctate actin signal. Instead, we observe that while traction forces require punctuated actin spots, not all actin punctae result in force generation. We call this a one-sided relation between in-plane tractions and punctate actin, which is accompanied by a homogeneously curved protrusion into the compliant gels. Correlating planar tractions and out-of-plane deformation with actin signal, we propose a scale-dependent spatiotemporal model of podosome cluster mechanics with buckling instability as the main driver for cluster protrusion.

## Results

### Surface Z plane extraction allows simple and fast 2.5D TFM and ventral actin projection

To gain deeper insights into the mechanics of cell protrusion, we considered a full capture of in-plane and out-of-plane tractions essential. While suitable and fast *free form deformation* (FFD) methods for computing 2D deformations in TFM are available ^26^, it gets more complicated upon adding the third dimension. We demanded a computationally efficient and fast algorithm to capture the dynamics of our multiplane image stacks. To this end, we developed an adapted algorithm implemented in *MATLAB* (MathWorks) for extraction of Z information from the respective bead stacks as illustrated in Figure 2. Briefly, we aimed at extracting the Z coordinates of the marker bead rich hydrogel surface to capture the literal ‘out-of-plane’ deformation as illustrated by the central, darker area (Fig. 2A). Hence, we calculated the bead stack’s locally grouped signal intensity along Z and defined the putative surface coordinate as the maximum of its B-spline fit (Fig. 2B). Repetitively interpolating the discrete values gained from this calculation generated a surface Z map that contained the gel surface’s Z coordinates. Correcting for the intrinsic gel and stage slopes we here achieved an 95 % uncertainty interval ranging from approximately -0.05 to +0.05 µm given a 0.5 µm Z plane distance from our experimental settings (Fig. 2C). The Z deformation field (Uz) could then be calculated by subtraction of the force loaded and unloaded surface Z maps, respectively. Access to 3D deformation fields (Ux, Uy, Uz) of the cell substrate enabled 2.5D TFM for the ER-Hoxb8 monocytes (Fig. 2D) in parallel with ‘2.5D imaging’ of the intracellular actin dynamics. Specifically, by application of the surface Z map’s coordinates to the Lifeact-EGFP channel it was possible to project the substrate-faced/ventral cell surface into a single plane for better actin signal/traction comparisons. It should be noted that this is neither a Z plane maximum projection, which would overlay the full actin signal of the cell, nor a single image plane. In contrast, this method specifically extracts the actin signal around the constructed curved plane above the deformed substrate.

**Fig. 2.**
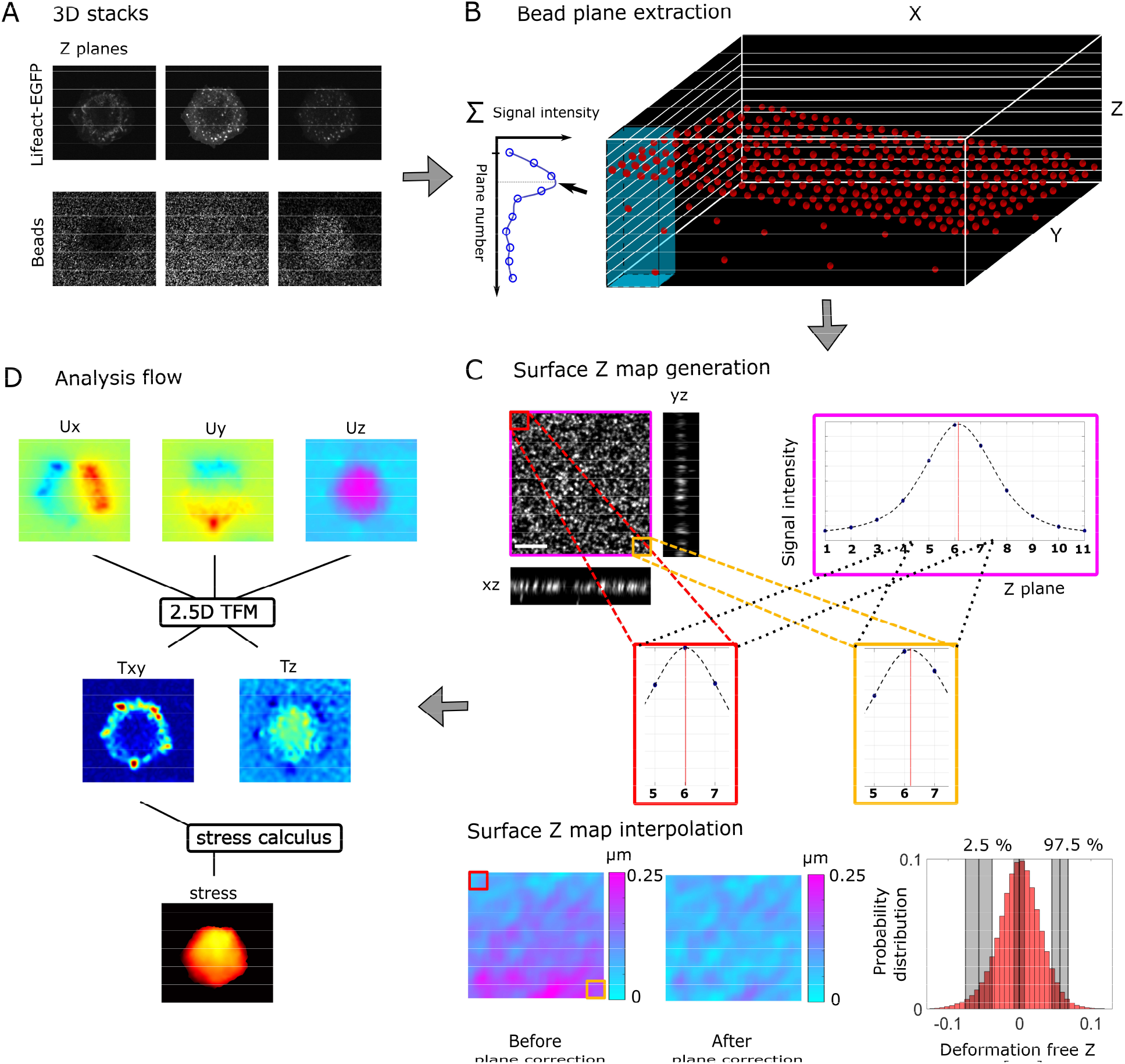
Workflow for out-of-plane/2.5D Traction Force Microscopy (TFM) **(A)** The Lifeact-EGFP and bead signals captured over several planes reveal punctate actin structures and a substantial Z deformation, respectively. **(B)** The bead planes‘ Z information can be extracted by an in-house MATLAB code that pillar-wise scans through the bead stack, **(C)** fits a B-spline to the intensity distribution along Z and takes the peak as the, true‘ local Z plane. Shifting over the entire bead stack and interpolating the discrete Z plane map yields a surface Z map of the gel (*bottom left*). The small but apparent intrinsic surface skew due to gel fabrication and mechanical stage slope is then corrected by subtracting a plane that is fit to the 10% border values of the Z map (*bottom center*). As a measure of uncertainty, we calculated the Z value distribution in the plane corrected areas of each gel that was used for shift correction (N=12) and normalized it by the number of frames (*bottom right*). The shaded areas indicate the standard distribution of the lower and upper limit of the 95% interval, respectively. Based on this, Uz-values of more than approximately 0.05 µm or less than -0.05 µm can be considered as surplus deformations. Consequently, the Z resolution for our experiments can be approximated to be in the range of 10-20% of the microscopy Z plane distance. Scale bar: 10 µm. **(D)** Applying the surface Z information to the Lifeact-EGFP and bead stacks enables us to extract the respective surface-faced signals. These information are then fed into our 2.5D TFM analysis to calculate the in-plane as wells as out-of-plane tractions. Based on the planar tractions, we then calculate the stress distribution due to the planar tractions using a FEM.

### Podosomes’ higher-order architecture is reflected in traction forces

We observed that Lifeact-EGFP labelled ER-Hoxb8 monocytes abundantly displayed dynamic podosome superstructures (Fig. 3A) upon treatment with S100A8/A9. Surprisingly, the cells presented with a large and dynamic morphological variety in podosome architecture 30 minutes after seeding onto the soft hydrogels. While most cells displayed cell wide podosome clusters, all three of the most abundant superstructures described in literature ^9,11^ were present. A cell demonstrating the dynamical transition of these superstructures on its ventral surface is shown in Figure 3A. The respective cell displayed ‘ring’, ‘belt’ and ‘cluster’ superstructures dynamically interconverting on the short time scale of 15 minutes. To our surprise, analyzing the respective tractions, each superstructure appeared with a distinct force pattern. The ‘ring’ presented with an outward directed pushing force as described previously ^27^. The ‘belt’ exhibited a rail-like traction pattern bordering its characteristic actin shape with an outward located pushing and an inward located pulling line, respectively (Supplementary Movie 1). A strong apparent correlation between actin signal and both Z deformation as well as planar traction patterns from ‘ring’ to ‘belt’ organization gradually vanishes as the podosomes scatter over the cell area in a cell wide cluster of single podosomes with an inward directed peripheral traction ring (Fig. 3A *bottom, shaded areas*). According to common models, actin is only able to generate contractile forces, as in a pushing case the filaments would buckle. Hence, the observation that the vector orientation changes from outward (‘ring’) to bidirectional (‘belt’) to inward directed (‘cluster’) suggests either a special relation, or a dependence on the actual podosome superstructure dynamics for the forces. Additionally, these outward directed forces suggest that at least part of the cortical actin network is sufficiently stiff to bear a certain degree of compression. Taken together, the temporal evolution of the traction force pattern and the disperse scattering of the actin signal into apparently individual podosomes suggests that a mesoscale effect is dominating over forces originating from individual podosomes.

**Fig. 3.**
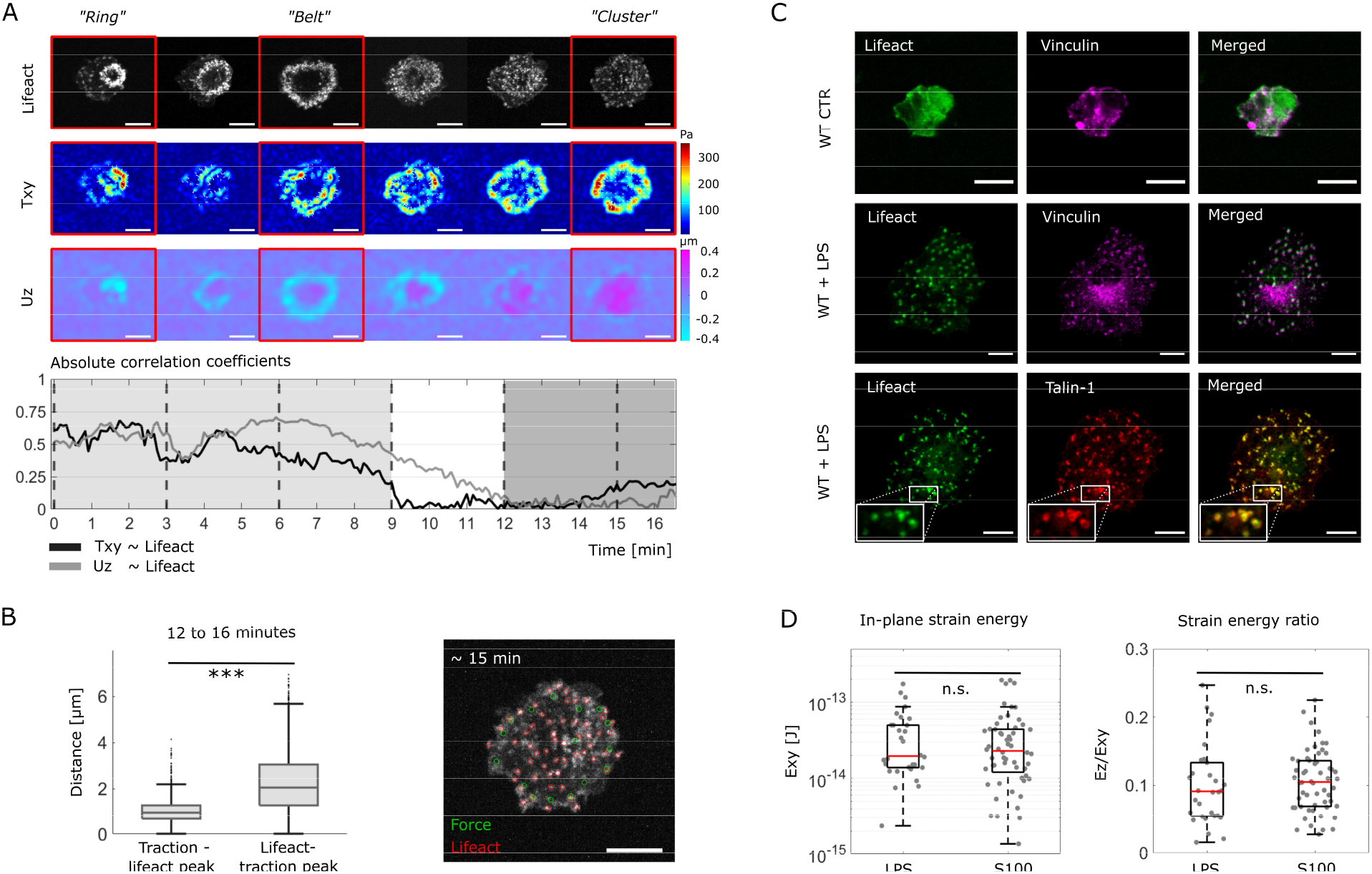
Podosome formation, dynamics and mechanical pattern in ER-Hoxb8 derived monocytes. **(A)** A selected ER-Hoxb8 monocyte treated with S100A8/A9 displays a rapid transition of three of the described podosome superstructures after seeding on a soft PAA hydrogel. Each superstructure presents with a distinct traction and Z deformation pattern. While from the, ring‘ to the, belt‘ structure a correlation between actin signal and mechanical pattern becomes apparent, during conversion into the most abundant, cluster‘, this relation disappears. Here, a circumscribed mechanical pattern is accompanied by diffusely spread actin cores. Projected, ventral Lifeact signal. Captured time interval: 5 seconds. Scale bar: 10 µm. **(B)** Though correlation disappears with appearance of the cluster, there still appears to be a one-sided relation between tractions peaks (green circles) and actin signal (red circles) regarding the mean distance (*left, p-value < 0*.*001*). **(C)** LPS-treated, Lifeact-EGFP labelled ER-Hoxb8 monocytes in contrast to untreated cells form punctate actin structures that colocalize with vinculin and talin-1, thus defining the observed structures as podosomes. A single plane of the respective image stack acquired by spinning disc microscopy is shown. Scale bar: 10 µm. **(D)** ER-Hoxb8 derived, podosome cluster bearing monocytes treated with LPS (n=30, N=3) or S100A8/A9 (n=56, N=5) display a similar mechanical behaviour in addition to actin morphology. In addition to similar median in-plane strain energies (*left, p-value 0*.*69*), the median ratio of out-of-plane (Ez) to in-plane strain (Exy) is not significantly different (*right, p-value* 0.32).

### Forces require podosomes, but not all podosomes generate forces

This finding encouraged us to investigate global mechanics in podosome bearing cells more thoroughly. The mesoscale mechanical effect was most apparent upon podosome cluster formation, which not only represented the most abundant phenotype in our study but was also widely investigated in previous macrophage and dendritic cell studies ^8,10^. Hence, we decided on focussing on the cluster superstructure for further analyses. The cluster with its mesoscale phenomenon was well characterized by a one-sided traction peak to punctate actin signal relation. This refers to the observation that traction peaks regularly appear close to actin punctae while the reverse is not valid upon podosome cluster formation (Fig. 3B). Although investigation of the effect of S100A8/A9 on leukocyte morphology and mechanics is a new, and hence attractive research direction, any generally relevant structure should also be observable using other activation approaches. To test if these observations are general for differently activated cells we moved to an exogenous alternative and treated our cells with *lipopolysaccharide* (LPS), which successfully induces both, a phenotype and a mechanical behaviour comparable to S100A8/A9 treatment (Fig. 3C+D). In the majority of cells being treated with LPS at a concentration of 1 µg/mL 24 hours prior to experiments ^28^, small punctate actin enrichments formed on the ventral surface and colocalized with vinculin and talin-1, in contrast to untreated cells (Fig. 3C). A direct comparison between LPS (n = 30, N=3) and S100A8/A9 (n = 56, N=5) treated, podosome cluster bearing ER-Hoxb8 monocytes showed a similar in-plane strain energy distribution as a measure of mechanical expenditure for both conditions. It roughly ranged in the magnitude of 10^−14^ to 10^−13^ Joule (p-value 0.69). Additionally, the median ratio of out-of-plane (Ez) to in-plane (Exy) strain energies that centered around 10 % was not significantly different (p-value 0.32) (Fig. 3D).

### Planar cluster tractions comprise two spatiotemporally distinct components

Triggered by this striking effect of cell stimulation and due to the absence of previous reports on the relation between traction generation via podosomes on the whole cell level, versus local forces at the single podosomes, we started with a general characterization of podosome cluster mechanics. In a first step we studied the phenomenological relation between podosomes and traction forces. Comparing localization of planar traction peaks and actin-rich punctae showed a regular occurrence of podosomes at sites of peak tractions (Supplementary Movie 2). Furthermore, while force peaks as mentioned before regularly corresponded to actin punctae, not all actin punctae corresponded to force peaks (Fig. 4A). Interestingly, the observed traction force centers were primarily located at the boundary of the cell (Fig. 4). The densely located traction peaks at the cell’s boundary accumulated to variously shaped rings and established as a common pattern throughout our analyzed cells.

**Fig. 4.**
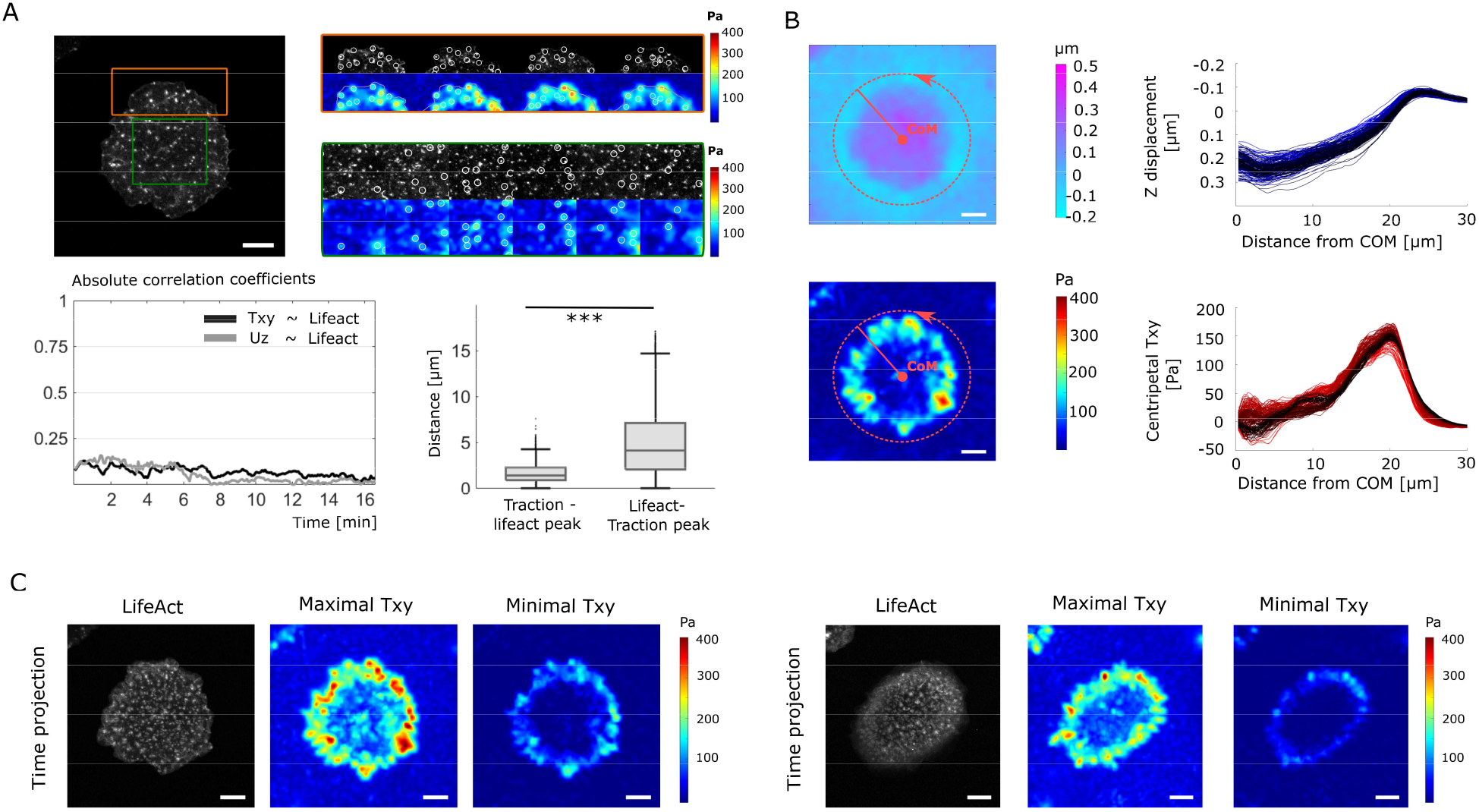
Mechanical pattern characterization in podosome forming ER-Hoxb8 monocytes. **(A)** The podosome cluster of a representative ER-Hoxb8 monocyte treated with LPS is evenly distributed over the cell’s ventral projection. While focal tractions are mainly aligned along the boundary (*orange*), some tractions appear centrally (*green*). Though without correlation, traction peaks are associated with podosomes but not vice versa (p-value < 0.001). Scale bar: 10 µm. **(B)** Radial profile (as illustrated by the *dotted red line*) of the Z deformation field (Uz) and the planar traction map (Txy) over time expose centripetal tractions close to the edge of the substrate indentation. In the centre, centrifugal (negative sign) and centripetal (positive sign) focal tractions arise. Later time points are indicated by darker colours. Scale bar: 10 µm. **(C)** Maximum time projections of the Lifeact-EGFP signal reveal high actin dynamics represented by podosomes covering almost the entire, variously shaped ventral area (*left* cell from A+B, *right* exemplary ellipsoid cell). In contrast, maximum and minimum time projections of the planar tractions expose transient central focal tractions and a stable peripheral traction zone. Scale bar: 10 µm.

In order to include space and time parameters into the analysis, we next characterized the spatiotemporal pattern of deformation and traction. To this end, we chose two simple analytical approaches: one to summarize the spatial mechanical pattern over time and the other to elucidate the temporal evolution by tracking the force centers. Regarding the spatial characteristics, we decided on a radial profile of the Z deformation and the planar force projected on the vector towards the cell’s center of mass (CoM). This effectively yields the force component acting towards the cell center as a function of distance from the cell center. Fig. 4B shows the median traction component towards the CoM (binned for the neighboured 5 pixel) and its local Z deformation, respectively. Positive traction components indicated centripetal, negative centrifugal orientation, while the smoothness of the curve referred to the degree of circular symmetry. As illustrated for the representative cell, the Z deformation presented as a rotationally symmetrical, relatively shallow, smoothly curved protrusion (Fig. 4B *top*) descending in the vicinity of a peripheral, centripetal traction ring (Fig. 4B *bottom*). Represented by darker colours for later time frames, a stronger planar traction comes along with a deeper indentation. The analysis shows that the indentation rapidly increases when seen from the edge of the cell, and slowly levels off towards the center (Fig 4B, Z-displacement). This suggests that the cell is not indenting the substrate most at positions with the highest centripetal Txy, hence lateral forces (Fig 4B). However, the change of substrate indentation is the highest at the edge, which we will quantify in more detail later as the slope of the indentation. While this method added directional information of the traction magnitude over time, local dynamics of the pattern were not being considered. To introduce dynamical aspects, we averaged over time by a simple time projection of the maximum values for the actin signal and the maximum as well as minimum values for the traction field (Txy), respectively (Fig. 4C). Surprisingly, the minimum and maximum traction projections taken together revealed two distinct traction components: a peripheral, in the observed time intervals spatially constant ring-like traction in addition to focal, transient tractions emerging over the entire cluster area. Opposingly, the maximum time projection of the actin signal revealed no correlative pattern but displayed a ventral cell surface with relatively dense and homogeneously scattered podosomes.

### High actin dynamics and two simple mechanical patterns characterize podosome clusters

Subsequent to the development of our tools for mechanical characterization we employed these to quantify vertical protrusion, hence indentation into the substrate (Uz) and planar tractions (Txy) in our podosome bearing cells (Fig. 5). Any forces, also traction forces inside cells have to obey Newton’s laws that state that the stress needs to be balanced in a steady state situation. In the context of the cell this means that the contractile stress that is generated on one side of the cell needs to be transmitted to the other side of the cell, to ensure conservation of momentum. In most other cell types, this is ensured by stress fibers spanning across the cell and the nucleus to connect the focal adhesions ^29^. However, in the absence of such stress fibers, other structures have to be in place to ensure force equilibrium inside the cell. To determine the predicted stress components, we again exploit Newton’s laws and use a finite element approach that allows to derive the stress field inside our cells, as previously demonstrated for other system ^30^. Representative traction force and stress results are shown in Figure 5A. Yet again, the five second acquisition interval exposes a highly dynamic punctate actin signal evenly distributed on the ventral cell surface (Fig. 5A *1*^*st*^ *row*). Opposingly, the mechanical patterns although displaying dynamics in magnitude do not show any substantial changes in topography. Instead, two distinct shapes characterizing the results stand out the most. First, a peripheral, circular pattern exhibited by the planar traction component (Fig. 5A *2*^*nd*^ *row*), which is reflected in the indentation slope (Fig. 5A 5^*th*^ *row*). Second, a central pattern demonstrating a homogeneous stress propagation within the cell, displayed as the internal normal stress (Fig. 5A *3*^*rd*^ *row*) and the absolute value of the indentation of the cell into the substrate that is here called the protrusion Uz (Fig. 5A *4*^*th*^ *row*). We confirm this visual correlation by the respective Pearson correlation between different pairs of quantities: Uz versus the internal normal stress, the central indentation (central Uz) versus the median centripetal tractions (Radial Txy) as well as the Uz slope versus Txy locally correlate with a Pearson coefficient of 0.96, 0.93 and 0.74, respectively. This quantitatively confirms the patterns’ interdependence as shown for the representative cell (Fig. 5B). Noteworthy, although the patterns show such clear interdependency, it is a non-trivial relation as the XY and Z based information are calculation independent.

**Fig. 5.**
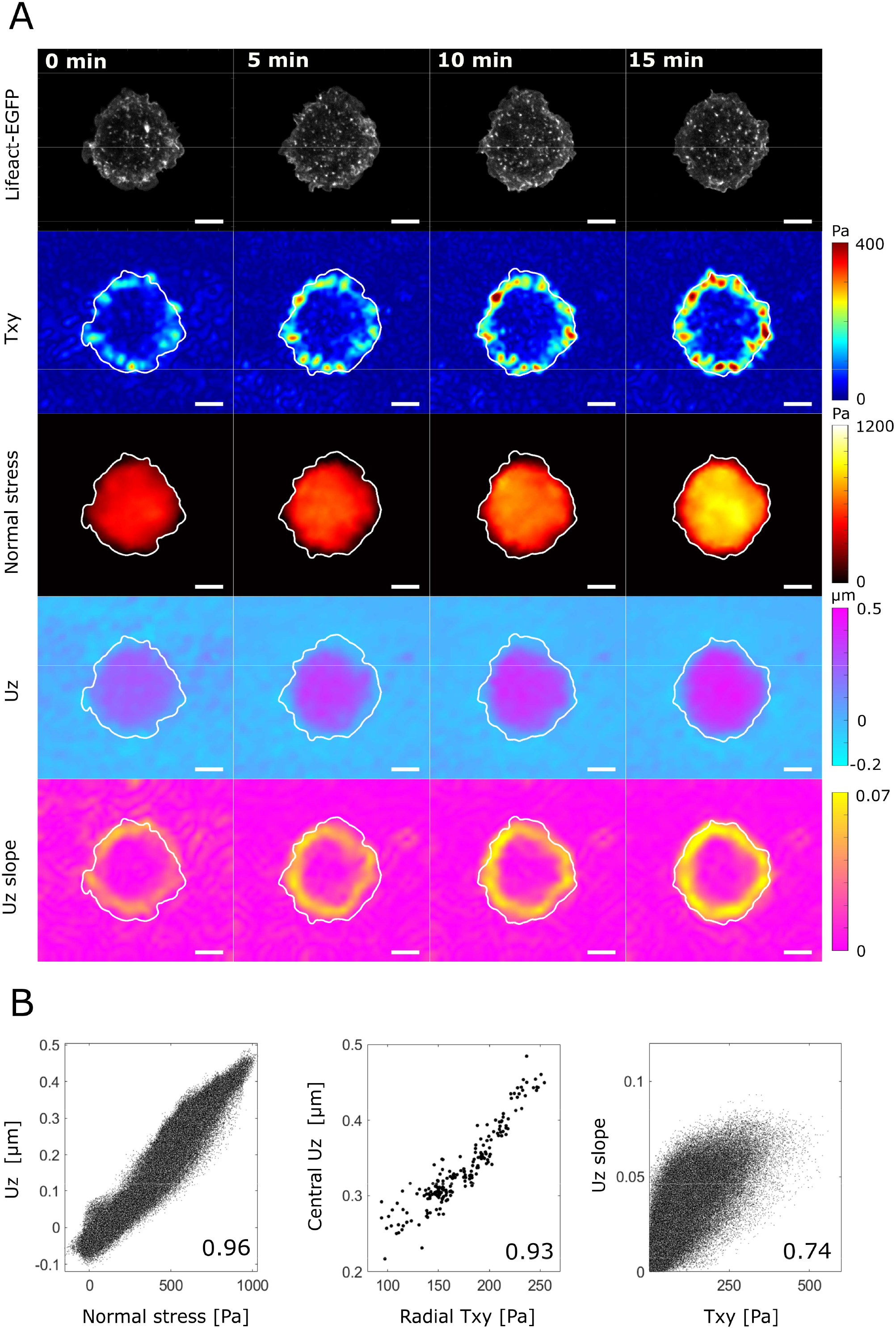
Exemplary mechanical analysis of a podosome forming monocytic cell. **(A)** The dynamic actin pattern (*1*^*st*^ *row*) is accompanied by a stable circular planar traction ring, that strengthens over time (*2*^*nd*^ *row*). The resulting stress builds up inside the circular area (*3*^*rd*^ *row*) associated with a similarly shaped out-of-plane deformation (4^*th*^ *row*). The Z deformation exposes a peripherally aligned peak slope (5^*th*^ *row*). Captured time interval: 5 seconds. Scale bar: 10 µm. **(B)** The internal normal stress due to Txy locally correlates with Uz (*left*, Pearson coefficient: 0.96). This is accompanied by a correlation between the median centripetal Txy and the central indentation yielded from the radial profile (*middle*, Pearson coefficient: 0.93). Last, the Uz slope moderately correlates with Txy (*middle*, Pearson coefficient: 0.93). For reasons of illustration, only every 250^th^ value is depicted in the first and third plot. LPS stimulation.

### In-plane and out-of-plane geometry favors a buckling induced protrusion behaviour

In general, the correlation between the intracellular stress (normal stress) that is transmitted through the cell and the out out-of-plane deformation can be achieved by either of two opposing mechanical mechanisms: redirected bending or buckling. Redirected bending as the ‘classical’ view considers the forces to be relayed from peripheral contact points (here supposedly podosomes) to the cortex or stress fibers and then intracellularly redirected onto the nucleus resulting in nuclear-driven pushing into the substrate. Hence, the intracellular stress is transmitted along the top of the cells, thereby pushing the intracellular material down. As only the nucleus has a reasonable stiffness, it will transmit these forces onto the substrate (Fig. 6A *left*). In this view the measured intracellular stress would not be transmitted at the ventral, hence substrate-faced part of the cell. In contrast, buckling considers the internal stress as located on the ventral plane and the vertical deformation as driven by and instability due to (here supposedly podosome) network contraction. In this view the internal stress transmission and the main substrate deformation are aligned perpendicularly (Fig. 6A *right*). While within certain limits both models are able to explain the correlation between internal stress and deformation, during force generation differences arise due to geometry. In the bending model substrate indentation (protrusion) occurs at the site of nuclear indentation. Consequently, the peak protrusion slope is supposed to be in the range of the nuclear indentation area and limited by nuclear size. In contrast, the buckling model predicts the highest slope to be close to or colocalizing with the underlying planar tractions. In our study we observe diversely sized and shaped peripheral traction patterns with only small differences between LPS and S100A8/A9 treated cluster bearing cells (Fig. 6B). The median radii as defined from the circular traction projection were 15 µm for LPS and 13 µm for S100A8/A9 treated cells (p-value 0.018) suggesting differences in either the overall cell size, or of the substrate interaction sites due to LPS and S100 activation. At these sites the two cell sets displayed widely scattered traction forces with no significant difference in their median traction that ranged at around 100 Pa (p-value 0.72) (Fig. 6C). Comparing the local Pearson correlation between the analyzed in-plane (Txy, stress) and out-of-plane information (Uz, Uz slope) exposed a strong correlation for internal normal stress and out-of-plane deformation (LPS: 0.90, S100A8/A9: 0.91, p-value 0.87) and a moderate correlation for planar tractions and local slope (LPS: 0.60, S100A8/A9: 0.52, p-value < 0.01) (Fig. 6D). Taken together, the buckling and not the bending model’s predictions were observed for both conditions, as we find a correlation between the indentation (Uz) and the internal stress. An important final prediction of the buckling instability is that the indentation will spontaneously occur once a critical lateral tension is generated. Indeed, we find this behavior when observing the time dependent stress built up. Upon the value of about 30 Pa has been reached, the cell starts to indent the substrate (Fig. 6E). This critical value is close to the predicted value of 75 Pa (see Material and Methods). In the classical indentation model, a linear dependence of the indentation on the tension in the cell would be predicted. Both, the observation of an instability and the good agreement with the prediction of the buckling stability supports that buckling is a process that should be considered when describing out of plane indentation generated by clusters of podosomes.

**Fig. 6.**
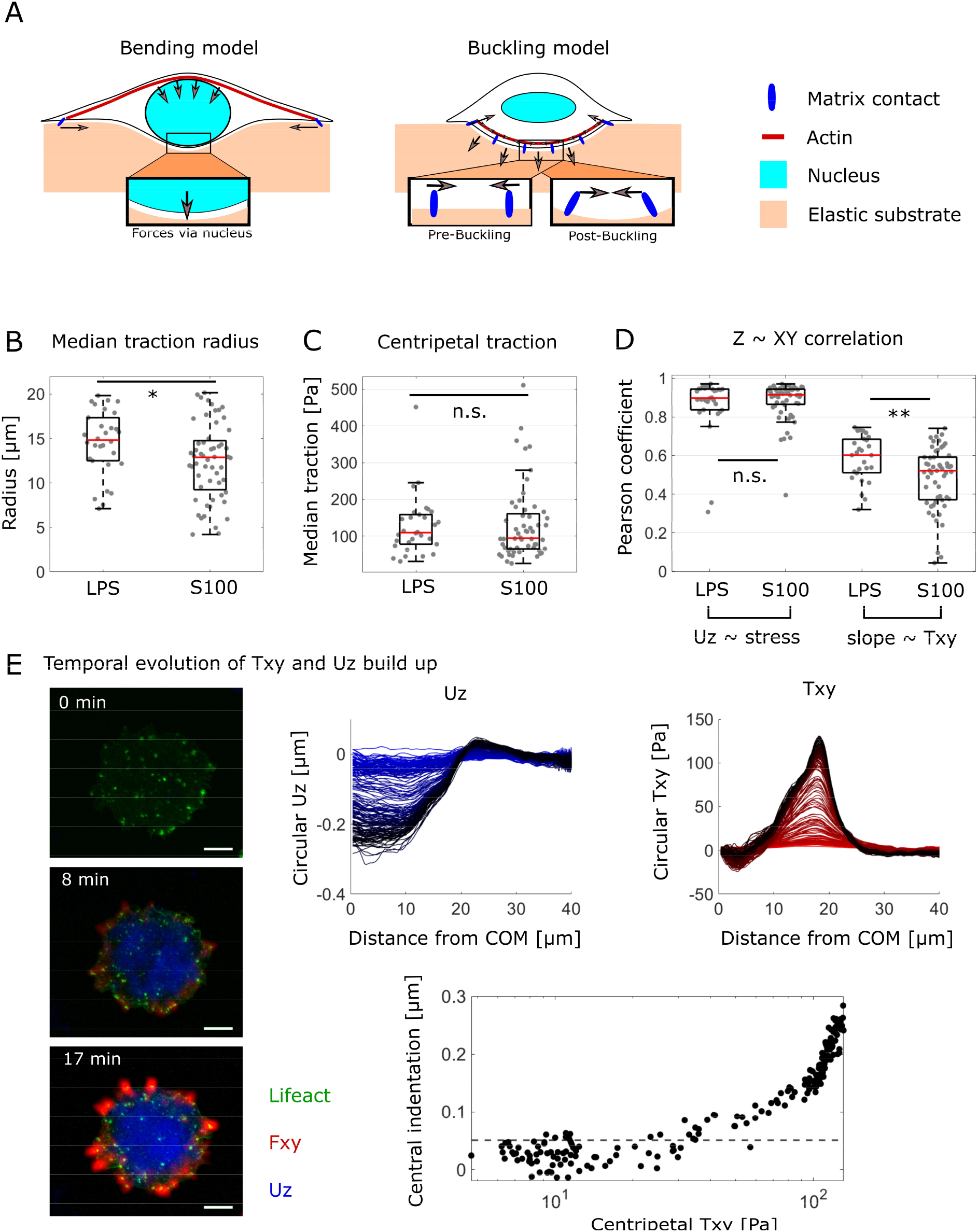
Two opposing mechanical models for out-of-plane traction generation tested against each other for plausibility. **(A)** The, classical‘ bending model (*left*) explains cellular out-of-plane deformation by substrate bending due to lateral focal adhesions that are linked to the cell cortex and/or the nucleus. They primarily exert upward directed tractions that are translated into downward directed tractions secondarily. The buckling model (*right*) describes substrate indentation by buckling due to contraction of a contractile network linked to the substrate. The network buckles if loaded above an instability threshold. **(B)** Both LPS and S100A8/A9 treated cells vary strongly in their median traction site radius. On average, S100A8/A9 traction radii are a little smaller (*p-value 0*.*018*). **(C)** The median traction at those sites does not significantly differ and ranges around 100 Pa (112 Pa vs 96 Pa, p-value 0.72). Comparing in-plane and out-of-plane mechanical properties reveals a strong median correlation between normal stress and Uz (0.90 vs 0.91, p-value 0.87) as well as a moderate median correlation between Txy and the Uz slope (0.60 vs 0.52, p-value < 0.01), respectively, supporting the geometrical buckling predictions. **(E)** An LPS treated cell exhibiting the rare event in which the cell displays tractions below and above the buckling threshold illustrates a steady sub-deformation (*dotted line*, 0.05 µm) range despite increasing planar tractions. Though without statistical weight it fits to the model’s prediction. Logarithmic X-scale. Scale bar: 10 µm.

## Discussion

Protrusion and degradation on the single podosome level have been widely investigated over the past years ^13,31–33^. New concepts were established linking degradation, actin structure, dynamics and force generation in the current model of podosome (mechano)biology during invasion ^11^. In parallel, the complex actin network of the podosome results in the unique formation of superstructures ^9^, which were deciphered to expose a multi-level organisation of F-actin with podosome-substrate and podosome-podosome connecting actomyosin ^13,16^. While mesoscale-connectivity and thus scale dependency was shown to play a role for dynamics ^8,12^ and putatively degradation ^3,34^, no such relation has been shown for mechanics. By employing 2.5D TFM, hydrogel surface-based protrusion and Lifeact-EGFP signal analyses on podosome cluster bearing monocytic cells, we here demonstrate a cluster boundary accentuated, in-plane traction and uniform central protrusion behaviour that do not recapitulate single podosome mechanics. Instead, they suggest being the mechanical correlate of podosome mesoscale-connectivity.

### Role of podosomes in global cluster mechanics

Analyzing the global mechanical patterns exposed by podosome clusters, we identified long-lasting, centripetal in-plane tractions at the clusters’ boundaries accompanied by transient, central, and focal tractions (Fig. 4C). Directly bordering to the peripheral tractions, we find a cluster filling shallow protrusion pattern that without further quantification was previously described ^18^. This mechanical behaviour entirely opposes single podosome mechanics which upon summation would suggest individual traction and protrusion spots even if passively coupled. As illustrated during temporal evolution of podosome superstructure transition, cluster formation initiates divergence of actin punctae and in-plane tractions, respectively (Fig. 3A). Noteworthy, a one-sided association remains between local peak tractions and actin punctae (Fig. 3B+4A). This suggests that podosomes likely keep being involved in force generation on the global scale but without each podosome spot transmitting these traction forces. In accordance with Newton’s laws, structures relaying these forces as internal stress are required. A structure present in various cell types fulfilling this function are actomyosin-based (ventral) stress fibers which span between focal adhesions ^29^. However, monocytic cells as fast migrating immune cells have not been reported to exhibit stable and long-lasting stress fibers or classical focal adhesions. Neither did we observe stress fibers in this study. Instead of large focal adhesions monocytes and macrophages present with small podosomes and instead of actomyosin-based ventral stress fibers linking focal adhesions they expose an interpodosomal actomyosin network. This network assumingly consists of layers coupled as well as uncoupled to cortical actin ^11,35^ suggesting a partially independent mechanical role from the actin cortex. As a result, everything required to generate and relay the observed in-plane tractions has already been reported to be present in podosome clusters.

### In-plane traction generation

Both the two-component traction pattern (Fig. 4) as well as the contractile, interconnected multi-element system show mechanistic parallels to epithelial monolayer mechanics ^30^. Both systems are characterized by mechanically active units that upon coupling via contractile actomyosin form larger aggregates with altered mechanics on the global scale. The similarities are further supported morphologically as the monolayer traction patterns and the generated stress strongly resemble our data (Fig. 5) ^36^. Based on this, we assume an analogous underlying mechanism of in-plane traction generation: The substrate connected podosomes experience omnidirectional myosin II driven contraction conveyed by the connecting cables and counterbalance each traction with oppositely directed traction to their neighbours. If entirely encircled by podosomes the net force becomes zero and no force is transmitted onto the substrate (Fig. 7A *right*) although the podosome finds itself under a high tension where all neighbours are pulling on it. If either the podosome is not entirely encircled by other podosomes, e.g. if located at the boundary, or if the omnidirectional connection is locally disrupted, forces are no longer internally relayed as stress but locally transmitted to the substrate (Fig. 7A *left*), which simply explains the high traction forces at the boundary of the cell.

**Fig. 7.**
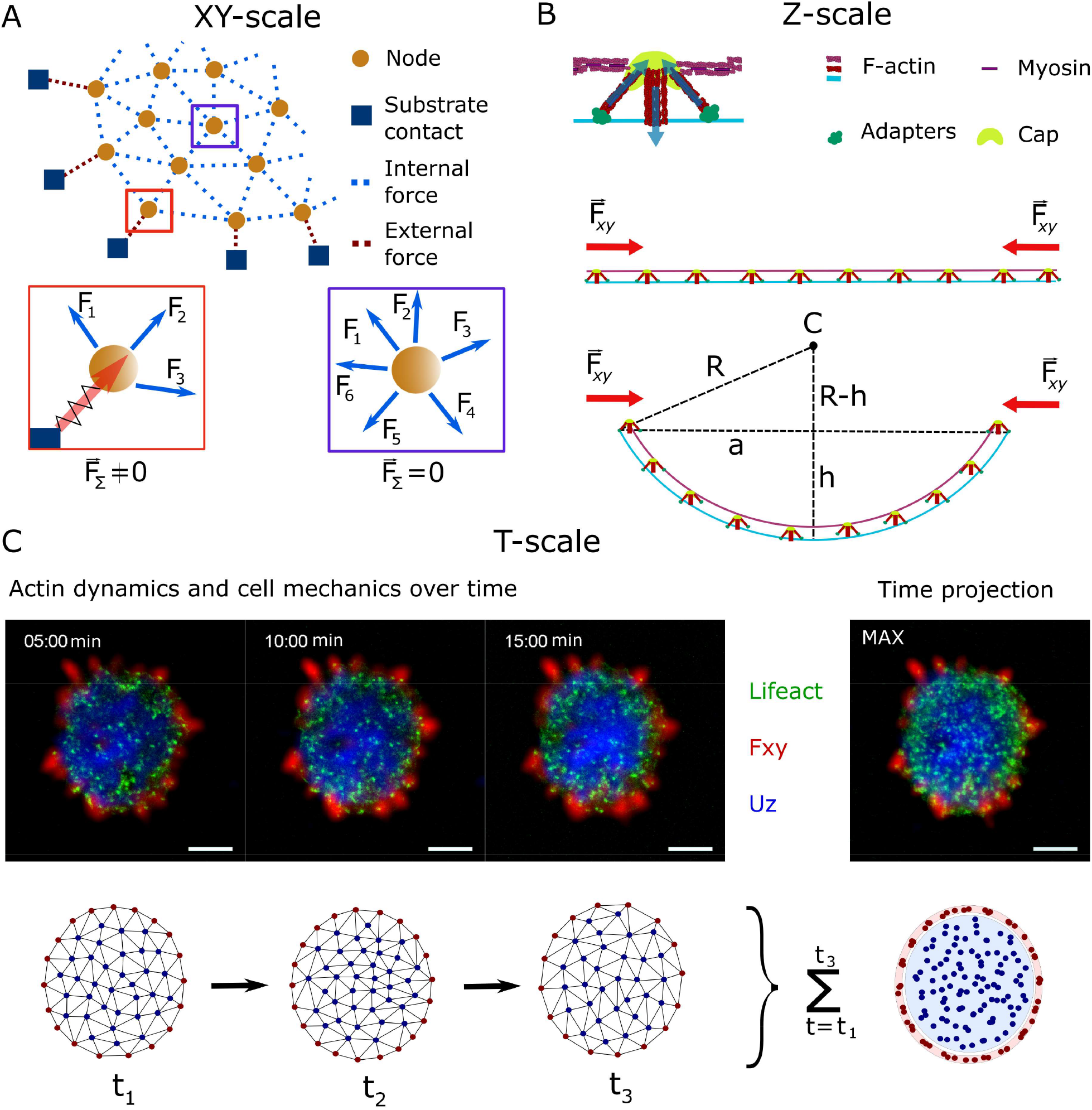
Proposed *XY-Z-T* model of podosome mesoscale mechanics. From our data we propose a model of podosome mechanics on the mesoscale in which in-plane (*XY*) tractions are generated by the prementioned interpodosomal actomyosin network. **(A)** Interpodosomal actomyosin generates contractile forces that are either relayed to neighbouring podosomes as internal stress (*blue dotted line*) or transmitted to the substrate as tractions by podosomes that are not omnilaterally coupled (*red dotted line*). **(B)** Upon contraction, the node-wise to the substrate coupled network builds up internal normal stress, starts to buckle upon reaching an instability threshold and protrudes into the substrate (*Z*). *R: curvature radius, C: curvature’s center, h: indentation, a: traction radius*. **(C)** As the generated tractions are exerted via the most peripheral podosomes that cannot relay forces omnidirectionally, the traction pattern appears stable at the edges (*red*) despite the changing node pattern covering the entire surface area over time (*T*) (*blue*). Given that the podosomes locally degrade their substrate, an almost cell covering degradation zone can arise in the area of protrusion (*blue*). Time interval: 5 seconds. Scale bar: 10 µm.

### Protrusion into the substrate

One difference between the mentioned epithelial monolayer and monocytes that is decisive for podosome cluster mechanics, is obviously its size. Epithelial cell monolayers are usually patterned to span over several hundred micrometers while the size spanned by the here observed clusters is in the order of 20 to 30 µm, hence much smaller (Fig. 6B). For in-plane traction generation this size difference becomes irrelevant because the stress is simply transmitted across the cell or the monolayer, respectively. However, out-of-plane mechanical effects, leading to substrate indentations, that usually are ignored in monolayer mechanics for size scale and proportion reasons ^37^ should not be ignored in our system. Instead, considering the contractile network that interconnects the podosomes inside the cells allows to speculate about an out-of-plane, hence pushing down, force that originates from an energy minimization due to a buckling like mechanism. While it is generally accepted that the origin for forces for both, the in-plane and out-of-plane is the actomyosin cytoskeleton, they are usually interpreted as local events occurring directly at the cell-matrix contacts ^38^. However, we simply do not observe that at the position of lateral traction forces (force spots at podosomes) we have the largest local indentation, but that we have a more global indentation that is correlated with the intracellular stress transmitted through the cell (Fig. 6D. left). Classical bending appears unlikely in the face of central, focal tractions, widely spanning and traction-associated peak slopes and the need to assume a not reported connection between cortical actin and substrate-attached integrins. However, we do know of a contractile actomyosin network that interconnects podosomes which enable substrate coupling via integrins. Hence, we propose a network contraction driven, instability induced buckling of podosome clusters. In the buckling picture, it is energetically more favourable to indent the substrate and therefore allowing to compact the contractile actin network, instead of only generating a 2D deformation of the substrate (see Materials and Methods). From the buckling a variously shaped, uniform, shallow protrusion pattern is able to arise (Fig. 7B). Noteworthy, we explicitly do not claim that single podosome mechanics are false but rather superimposed on the mesoscale as illustrated in Figure 3A. In our model scale dependent podosome mechanics relates single to collective mechanics by a *musketeer effect* (*‘one for all and all for one’*): While single podosomes connect to the substrate and locally degrade the substrate, on the larger scale the collective mediates network buckling induced protrusion and thus allows the single podosomes to keep contact to the gradually degraded substrate. Hence, not all podosomes are required to be substrate-connected at each timepoint.

### Actin dynamics and deducible predictions for degradation

This behaviour is further supported by the apparently stochastic process of podosome scattering as a result of high dynamics and short lifetimes as illustrated in the time projections (Fig. 4C). Taking into account that degradation mediating MT1-MMP expression was reported to colocalize with emerging podosomes ^33^, the time projections indicate a homogenous degradation pattern as reported before ^3^ instead of a dotted degradation pattern similar to invadopodia ^18^. Since the widespread degradation of matrix proteins would render almost all individual podosomes nonadhesive and thus tractionless, single podosomes would be unable to convey protrusion through the formed gap. However, our data provide a solution to this dilemma as both traction transmission and cluster wide protrusion are spatially separated and thus remain achievable on the mesoscale.

### Model system caveats

Since the model of ER-Hoxb8 cells is relatively new to the field it was not extensively tested for podosome properties yet. Even more importantly, some apparently incoherent results to literature might arise as our ER-Hoxb8 cells differentiate to macrophages by protocol while many studies have investigated dendritic cells which have shown to regularly behave substantially different, e.g. upon TLR4 stimulation and integrin expression ^39,40^. Hence, it is not surprising that LPS does not induce podosome cluster formation in other cells although for ER-Hoxb8 monocytes the effect appeared to be reproducible. The second caveat arises from our TFM method: While we decided for a better temporal than spatial resolution, although sufficient for the mesoscale, we were not able to resolve the previously reported single podosome tractions and merely assume that these are superimposed to our findings. Meanwhile, while providing some evidence we are unable to show the proposed mechanical hierarchy from our data alone and are required to refer to previous papers as support for our model. Finally, we might overinterpret the podosomes’ and in particular the actin’s function for mesoscale mechanics and traction generation simply by employing Lifeact-EGFP labelled cells while ignoring all other proteins involved in this complex process.

To this end, studies to further characterize the mechanical hierarchy by improved spatial resolution as well as simultaneous quantification of cell wide degradation and protrusion as shown for invadopodia ^17^ are required to gain deeper understanding in the mechanism of podosome facilitated barrier breaching. Additionally, while we finally focussed on podosome clusters, we have provided evidence that podosome superstructures come with their respective and distinct mesoscale mechanics. Resolving connectivity and mechanics here might gain deeper insights into the respective biological roles of these higher-order structures and the reason for transition.

## Conclusion

In conclusion, in this study global tractions and protrusion of podosome clusters were characterized. We show that on the podosome cluster scale the spatially resolved protrusions are consistent with previously reported degradation sites that follow the clusters’ shapes ^34^ and thus mechanistically enable barrier breaching on the cell scale. By resolving a two-component, boundary accentuated traction that cannot be delineated by single podosome mechanics we demonstrate a mechanical correlate of podosome mesoscale-connectivity. Based on this, we propose a scale dependent mechanical model including the interpodosomal actomyosin network. In this spatiotemporal mesoscale model myosin driven network contractility (*XY*) generates planar tractions while building up internal stress (Fig. 7A). Upon reaching an instability threshold, the network buckles into the substrate resulting in cluster wide protrusion (*Z*) (Fig. 7B). As the network mechanics are supposed to originate from nodes being connected to the substrate but not their localization, single podosome’s rapid (re)emergence does not substantially affect the network’s mechanical pattern as long as coupling to the substrate is maintained (*T*) (Fig. 7C).

## Methods

### Generation of ER-Hoxb8 cells (monocytic lineage)

Bone marrow cells of Lifeact-EGFP mice were isolated from femur and tibia by flushing with PBS containing 1 % FCS (PBS/FCS). After lysis of erythrocytes and washing in (PBS/FCS), cells were differentiated for up to 7 days in RPMI1640 medium supplemented with 10 % FCS, 1 % Pen/Strep, 1 % L-Glutamine and 20 % supernatant of M-CSF expressing L-929 cells (Lonza).

Generation of Hoxb8 cell lines were made as previously described ^20^.

### Cell culture of ER-Hoxb8 cells

Cell culture and differentiation was according to recent work ^28^ with minor adaptations. Briefly, ER-Hoxb8 cells were cultured in uncoated six-well culture plates in RPMI 1640 medium (Merck Millipore) supplemented with 10% FBS (Pan Biotech), 1% L-Glutamin (Biochrom), 1% Penicillin/Streptomycin (Biochrom), 2% GM-CSF (ImmunoTools, Friesoythe, Germany), and 1 µM β-estradiol (Sigma-Aldrich, Steinheim, Germany). Cells were split every 2 days.

### Differentiation and stimulation of ER-Hoxb8 derived monocytes

ER-Hoxb8 cell suspension was washed and centrifuged two times with PBS/10% FBS to remove β-estradiol. Subsequently, 0.5 - 1 × 10^6 Lifeact-EGFP labelled ER-Hoxb8 cells were seeded in 3 ml RPMI 1640 medium supplemented with 10% FBS, 1% L-Glutamin, 1% Penicillin/Streptomycin and 20% supernatant of M-CSF expressing L-929 cells (Lonza) in untreated six-well culture plates. Cells were differentiated for 3 days. Non-adherent cells were aspirated and discarded after 2 days. To detach the adherent monocytes, PBS supplemented with 2 mM EDTA (Roth, Karlsruhe, Germany) was used.

For stimulation, differentiation medium supplemented with S100A8/A9 (homemade, Münster, Germany) or LPS (L4391, Sigma-Aldrich, Steinheim, Germany) at a final concentration of 1 µg/mL was added to differentiating cells. Cells were exposed to S100A8/A9 for up to 48 hours prior to experiments and displayed punctate actin clusters. Incubation with LPS was limited to 24 h reducing toxicity while inducing a comparable phenotype.

### Traction force experiments

On day 3, towards monocytes differentiated ER-Hoxb8 cells were centrifuged second to detachment and resuspended in their previous differentiation medium. Subsequently, cells were seeded at numbers of 20 000 to 25 000 cells per gel (12 mm diameter) and incubated for 30 minutes. Having removed non-adherent cells, 2 mL differentiation medium supplemented with 25 mM HEPES (Millipore, Burlington, USA) were added. Within the next 1 to 2 hours, 4D stacks were captured at 5 second intervals and 0.5 µm Z plane distance employing a spinning disk system (CSU-W1 Yokogawa) in combination with a heating chamber set at 37 °C. Podosome cluster bearing cells included into the study were morphologically defined as displaying scattered actin punctae that did not align into belts or rings.

### Gel preparation

If not otherwise stated all chemicals were purchased from Sigma-Aldrich (Merck). Polyacrylamide (PAA) gels were prepared as previously described ^41^ with some modifications. First, glass bottom dishes (CELLview 35/10 mm, Greiner Bio-one International) were cleaned with 70% ethanol followed by 0.1 N *NaOH*. Afterwards, the glass bottom was covered with 200 µL *(3-Aminopropyl) trimethoxysilane* (APTMS) for 3 minutes, thoroughly washed and covered with 500 µL 0.5% glutaraldehyde for 30 minutes. In the meantime, PAA gel premix was prepared by adding 4 µL of *acrylic acid* to 250 µL of 2% *N,N’-Methylenbisacrylamide* and 500 µL of 40% µL *acrylamide* solution. For 3 kPa stiff PAA gels 75 µL of this solution were gently mixed with 415 µL of 65% PBS and 10 µL of fluorescent bead solution (100 nm, NH2 coated micromer-redF, Micromod). Polymerization was induced by 5 µL of 10% *ammonium persulfate* solution (APS) (Roth) and 1.5 µL *N,N,N’,N’-Tetramethylenediamine* (TEMED) (Sigma Aldrich) (Roth). Functionalization for coating was performed by activation of the acrylic acid with 0.2 M *N-(3-Dimethylaminopropyl)-N′-ethylcarbodiimide hydrochloride* (EDC), 0.1 M *N-Hydroxysuccinimide* (NHS), 0.1 M *2-(N-Morpholino) ethanesulfonic acid* (MES) and 0.5 M *NaCl* for 15 minutes at room temperature, followed by thorough washing with PBS and incubation with fibronectin (Sigma Aldrich) (50 μg/ml) and incubation at 37°C for 1 h or at 4°C overnight. The gel’s stiffness was confirmed by rheological measurements (AFM) to be consistent with the protocol.

### Z plane extraction from Bead planes

For Z plane extraction of the hydrogel’s surface, the framewise bead channel stacks (11 planes, 0.5 µm plane distance) were processed in MATLAB (R2020a, MathWorks Inc., MA, USA). Briefly, we here used columnar shaped (21 × 21 pixel) substacks which shifted over the gel area at a predefined overlap (15 pixel). From these, we repetitively performed a cubic spline interpolation to the signal intensity values along Z. For interpolation the increment number was increased by the factor 100. The spline fit’s peak was calculated and the value then assigned to the column’s center coordinate to fill a discrete matrix prior to cubic interpolation. In order to project the surface-faced Lifeact-EGFP signal, the surface Z map was applied to the respective stack to locally project the maximum from one plane beneath to one above the extracted plane number. For the accompanying drift correction we used ‘dftregistration’ ^42^ and as a template randomly chose a 400 × 400 pixel sized cell-free submatrix.

### Traction force and stress calculation

For traction force calculation, the in-plane deformation of the gel was derived for all timepoints from the deviation of the localizations of the fluorescent beads inside the gel at the respective timepoint from the bead localizations after cell detachment by applying a free form deformation (FFD) algorithm using the software Elastix ^26^. The calculation was carried out in a three-level pyramid approach from a coarse to fine scale. Grid size was divided in half and number of iterations doubled for each level. The grid size for the finest scale was set to 1.3 × 1.3 µm or 1.5 × 1.5 µm if generating excessive artefacts. Cells that displayed artefacts even with the larger grid were excluded from the study (LPS: n=1, S100A8/A9: n=2). The final result was reached optimizing the advanced Mattes mutual information metric using the adaptive stochastic gradient descent algorithm for 2000 iterations at the finest scale. The number of random spatial samples per iteration was chosen based on the size of the respective image. The out-of-plane deformation was calculated from the reference surface Z map which was extracted as described before being subtracted from the force loaded surface Z map. Prior to inserting into the traction calculation algorithm it was undeformed inversely applying the in-plane deformation fields.

The three dimensional traction forces exerted by the cells on the gel surface were then inferred from the displacement fields solving the Tikhonov regularized equation of the elasticity problem for finite thickness substrates in the Fourier domain ^43,44^ using a custom-made MATLAB program. To achieve a less subjective and more stable choice of the regularization parameter, Tikhonov regularization was carried out applying Bayesian theory combined with an estimation of the background variance of the deformation field as recently proposed ^45^.

Evaluation of strain energies was restricted to the cell area as outlined by the fluorescence signal of each individual cell. Cell mask binarization was performed employing the MATLAB implemented ‘activecontour’ function. Before elementwise multiplying traction and deformation along X, Y and Z, the respective cell free area’s median was subtracted as a background noise reduction. This is important as the energy noise accumulates in an area dependent manner.

Internal stress calculation from the in-plane tractions was performed using the COMSOL Multiphysics software package with LiveLink for MATLAB. Based on the aforementioned area representing the cell outline, extruding the latter into 3D with a height of 5 µm, a cell object was created modelling the cell as a homogenous, linear elastic material. Assuming force balance the formerly calculated traction forces were imposed as boundary load (with opposite sign) to the bottom plane of the cell object. A finite element mesh for the object was created using the automatically designed tetrahedral mesh with size setting ‘finer’. Further, the boundary condition ‘rigid motion suppression’ was applied and the material set to be nearly incompressible. Since small rotational artifacts could occur around an initial orientation dependent point, the calculation was carried out for four different in-plane orientations (each orientation rotated by 90°) of the cell object. Omitting the artificial regions for each rotation, the final result was derived as an average of all rotations. From the solution of the finite element equations the in-plane stress tensor at the bottom plane of the cell object was extracted. The internal stress presented in the results is given as the average normal, principal stress at each point, i.e. the mean of the xx- and yy-component of the stress tensor. For faster computation times the resolution of the traction force input data was reduced by a factor of 10 (object size was accordingly reduced for calculation) and by that naturally also the resolution of the resulting internal stress output.

### Slope calculation

As a measure of Uz curvature, we quantified the slope at each coordinate. To this end, we resized the Uz map to its original 21 pixel resolution and adapted the unit to pixel length. From the gradient fields along × and Y, respectively, we geometrically derived the dimensionless XY surface slope. Resizing back yielded a map corresponding to the fluorescent image as well as traction, Uz and stress fields for comparison. Due to limitations in Z resolution, time frames with an indentation depth of less than 0.1 µm were excluded for the Txy-slope correlation calculation (Fig. 6D).

### Actin and traction peak distance calculation

Local actin peaks were detected from the Lifeact-EGFP fluorescence channel in a three-step process of noise reduction, binarization and object filtering. Briefly, the ventral cell projections were 3D median filtered (kernel size: 5×5×5) followed by performing morphological operations to reduce irregularities. Local peaks were then identified employing the built-in function ‘imregionalmax’. The detected objects were further filtered by shape (‘regionprops’: ’Eccentricity’) as well as local absolute and relative intensities. Regarding shape and absolute intensity, the thresholds were finely modulated for each cell in order to best exclude membrane artefacts. This is essential as the traction pattern is also located at the cell’s boundary. If not excluded properly, it results in a false actin-traction colocalization. To this end, we used a conservative approach to rather ignore valid podosome peaks inside than to include actin-based membrane fluctuations at the periphery as erroneous peaks. For the given examples, eccentricity was set 0.3 for the selected cell in Figure 3A and 0.45 for Figure 4A while the absolute intensities were defined by the 95th and 97.5th percentile of the cell’s intensity values, respectively. Intensity filtering was performed based on the detected peaks’ 3×3 pixel environment. Regarding the relative intensity, we defined the local background’s intensity as the coarsely median filtered (kernel size: 50×50) image. and calculated a factor by which valid peaks had to surpass this local background. This factor was calculated from the 70th percentile of all intensity values inside the cell’s boundary that were greater than 110% the local background.

Traction peaks were detected similarly. Here, of all traction values inside the cell’s boundary that were greater than two times the 90th percentile of the cell free background, the 70th percentile was defined as the force threshold. Due to resolution, we only considered peaks that were more than 10 pixels (∼1 µm) apart.

Finally, as a measure of colocalization, we computed the distance between actin punctae and traction peaks by a MATLAB-implemented k-nearest neighbor search (‘knnsearch’).

### Buckling instability

To establish if the forces generated in the coupled system of cells to the substrate are sufficient for the proposed buckling, we use a classical energy approach comparing the energy required for the buckled situation with the energy needed in a simple 2D compression. First, we establish the energy contributions for bending the cell cortex, indenting the substrate and compressing the cell cortex. Estimating the contribution shows that the bending of the cortex is negligible compared to the indentation of the substrate. Balancing the energy contribution leads to a critical traction force at which buckling is expected.

Energy to bend the actin cortex: The bending energy of a thick layer is defined as 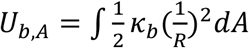, where, *k*_*b*_ is the bending modulus and *R* is the radius of curvature for the bending. Simple geometry shows that 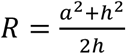, where *a* is the radius of the cell and *h* is the indentation from the unbend situation (Fig. 7b). Assuming a constant curvature and the limit of *a≫h* the integral simply yields the area of the cell resulting in:

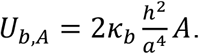

This energy is found to be small compared to substrate deformation energy which can be derived using the Hertz solution for the force to indent an infinite half plane with a sphere: 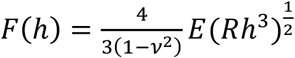, where *E, v* are the substrate’s Young’s modulus and Poisson’s ratio, respectively. Approximating the Poisson’s ratio with 0.5 and integrating the force over the indentation, keeping in mind that 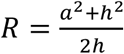, where a≫h, yields the deformation energy as 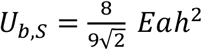

The energy to compress the actin cortex is simply given by *U*_*c*_ = *κ*_*A*_ × *ΔA* using the compression modulus *κ*_*A*_.

Using known values of these parameters either from literature, or as measured here we see that the bending of the cell cortex is negligible compared to energy required for indenting the substrate. The explicit values used are: *κ*_*b*_ = *κ*_*A*_*t*^2^, where *t* = 100 *nm* is the thickness of the actin cortex, and *κ*_*A*_ = *E*_*a*_*t* can be calculated from a stiffness *E*_*a*_ ≈ 1 *kPa* of the actin cortex. *h* ≈ 0.25 µ*m, a* ≈ 5 µ*m* and *A* = *πa*^2^ describe the geometry of the cell and *E* = 3 *kPa* is the Young’s modulus of the substrate.

The buckling will then happen at a critical traction stress. Beyond this critical stress, it is more energy efficient to buckle than to compress the actin network. This critical tension can be calculated by equating the bending and the compressing energy and keeping in mind that the area change can be calculated using the conservative approximation that the traction force is fully applied on the actin cortex *σ* = *E*_*a*_*u*, where *u* is the strain that is approximated to the relative area change in the compression situation. Hence the buckling condition is: *U*_*c*_ > *U*_*b*_, which yields 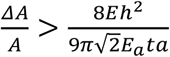, leading to the critical traction force for the given parameters of 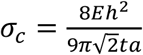. In the given situation we would hence expect the system to buckle if the traction forces are larger than 75Pa, which we are exceeding even with the median forces of 100 Pa. It should be noted that we here used conservative estimates, meaning that the real buckling might happen already much earlier.

### Immunofluorescence and microscopy

LPS-treated or untreated Lifeact-EGFP ER-Hoxb8 derived monocytes (day 3) were seeded onto 3 kPa PAA gels at numbers of 25 000 and incubated for 1 h. The cells were then washed once with calcium and magnesium free PBS (PBS^-/-^) and fixed in 4 % paraformaldehyde (PFA) for 15 minutes at room temperature. Afterwards, the sample was washed three times with PBS^-/-^ again and blocked for 1 h at room temperature using PBS^-/-^ supplemented with 20 % goat serum (GS, Sigma, St. Louis, USA) and 0.2 %Triton-X-100 (Carl Roth, Karlsruhe, Germany). For mouse-on-mouse staining, the cells were blocked for 1 h at room temperature using an unconjugated goat anti-mouse IgG H&L antibody (polyclonal, 1:1000, Abcam ab6708, Cambridge, UK) diluted in previously mentioned blocking solution. Next, the samples were incubated with the primary antibody (monoclonal mouse anti-vinculin, 1:100, Sigma V9264, St. Louis, USA; polyclonal rabbit anti-talin 1, 1:100, Abcam ab71333, Cambridge, UK) diluted in blocking solution for 1.5 h at room temperature or overnight at 4 °C.

After three washes with PBS^-/-^, the samples were incubated with the appropriate secondary antibody (polyclonal goat anti-mouse IgG1, 1:500, ThermoFisher A21124; polyclonal goat anti-rabbit IgG H&L, 1:500, ThermoFisher A11011, Waltham, USA) diluted in blocking solution for 45 min at room temperature. F-actin was stained using Phalloidin (1:1000, Abcam ab176753, Cambridge, UK). Finally, the samples were washed three times with PBS^-/-^.

Confocal images were acquired using the Slidebook 6 software (3i, Denver, USA) using an inverted microscope (Nikon Eclipse Ti-E, Minato, Japan) equipped with a CSU-W1 spinning disk head (Yokogawa, Musashino, Japan) and a scientific CMOS camera (Prime BSI, Photometrics, Tucson, USA). Images were analysed and prepared for publication using the open source software Fiji ^46^.

For TFM experiments images were acquired with a scientific CMOS camera (Orca-flash4.0 v2; Hamamatsu Photonics K.K.) on an inverted microscope (Nikon Eclipse Ti-E, Minato, Japan) with a spinning disk head (CSU-W1 Yokogawa, Musashino, Japan). A 60X Plan Apo water-immersion objective with a numerical aperture of 1.2 and excitation LASER of 488 nm/561 nm wavelength were employed.

### Statistical analysis

Statistical significance was tested against using the MATLAB-implemented Mann-Whitney U-test (‘ranksum’). The significance level α was set to 0.05. P-values < 0.05 are indicated by *, p-values < 0.01 by ** and p-values < 0.001 by *** throughout the study. The boxes of the presented box plots indicate the median in addition to the 1st to 3rd interquartile range. Pearson correlation coefficients were calculated using the MATLAB-implemented ‘corr’ function.

## Data availability

The data supporting our findings are available from the authors upon reasonable request.

## Code availability

The code utilized in this study is available from the authors upon reasonable request.

## Author contributions

TB and HS designed the study. HS, AR and AH performed experiments. MB, TB and HS wrote the MATLAB scripts. TB and HS created the figures. TB and HS developed the model. TV and JR contributed the model system. TB and HS wrote the manuscript. AR, AH, MB, TV and JR contributed to the manuscript. All authors read the manuscript. TB supervised the project.

## Acknowledgements

We thank Roland Wedlich-Söldner for kindly providing the anti-talin 1 antibody and Heike Hater for her support in ER-Hoxb8 cell cultivation. HS was supported by the MedK Program of the University Münster and by the SFB 1009, Breaking Barriers. TB was supported by the “Interdisziplinäres Zentrum für Klinische Forschung” (IZKF, Bet1/013/17) of the medical faculty, University of Münster, and by the ERC-Consolidator grant PolarizeMe (771201).

## Notes

### Competing Interest Statement

The authors have declared no competing interest.

