## Supplementary material for "2.5D Tractions in monocytes reveal mesoscale mechanics of podosomes during substrate indenting cell protrusion": Suppl. information

### Supplementary information

**Movie1: Relation of podosome (super)structure and mechanical pattern.** S100A8/A9 stimulated cell in Fig. 3A. From left to right: Lifeact-EGFP signal, planar tractions, Z deformation. Colourmap limits from Fig. 3A. Time format mm:ss. Scale bar: 10 µm.

**Movie2: One-sided relation between planar tractions and Lifeact-EGFP punctae.** LPS stimulated cell in Fig. 4A. Lifeact-EGFP signal (*green*) and planar tractions (*red*). Time format mm:ss. Scale bar: 10 µm.
